# Phylogenomics of the olive tree (*Olea europaea*) disentangles ancient allo- and autopolyploidizations in Lamiales

**DOI:** 10.1101/163063

**Authors:** Irene Julca, Marina Marcet-Houben, Pablo Vargas, Toni Gabaldón

## Abstract

**Background:** Polyploidization is one of the major evolutionary processes that shape eukaryotic genomes, being particularly common in plants. Polyploids can arise through direct genome doubling within a species (autopolyploidization) or through the merging of genomes from distinct species after hybridization (allopolyploidization). The relative contribution of either mechanism in plant evolution is debated. Here we used phylogenomics to dissect the tempo and mode of duplications in the genome of the olive tree (*Olea europaea*), one of the first domesticated Mediterranean fruit trees.

**Results:** Our results depict a complex scenario involving at least three past polyploidization events, of which two -at the bases of the family Oleaceae and the tribe Oleeae, respectively- are likely to be the result of ancient allopolyploidization. A more recent polyploidization involves specifically the olive tree and relatives.

**Conclusion:** Our results show the power of phylogenomics to distinguish between allo- and auto- polyplodization events and clarify the conundrum of past duplications in the olive tree lineage.

## Background

The duplication of the entire genetic complement -a process known as polyploidization or whole genome duplication (WGD)- is among the most drastic events that can shape eukaryotic genomes[1]. Polyploidization can be a trigger for speciation[2], and can result in major phenotypic changes driving adaptation[3]. This phenomenon is particularly relevant in plants, where it is considered a key speciation mechanism[4,5], and where the list of described polyploidizations grows in parallel with the sequencing of new genomes[6–11]. Polyploidization in plants has been a common source of genetic diversity and evolutionary novelty, and is in part responsible for variations in gene content among species[3,4,12].Importantly, this process seems to have provided plants with traits that make them prone to domestication[13], and many major crop species, including wheat, maize or potato are polyploids[6,10,14,15].

Polyploidization can take place through two main mechanisms: namely autopolyploidization and allopolyploidization. Autopolyploidization is the doubling of a genome within a species, and thus, resulting polyploids initially carry nearly-identical copies of the same genome. Allopolyploids, also known as polyploid hybrids, result from the fusion of the genomic complements from two different species followed by genome doubling. This genome duplication following hybridization enables proper pairing between homologous chromosomes and restores fertility[16–18]. Such mechanism has been described as the fastest (one generation) and most pervasive speciation process in plants[19,20]. Hence, allopolyploids harbor chimeric genomes from the start, with divergences reflecting that of the crossed species.

Elucidating the exact number and type of past polyploidization events from extant genomes is challenging. In part because following polyploidization a process called diploidization sets in, during which the genome progressively returns to a diploid state[4,21]. This is attained through massive loss of genes and even of whole chromosomes, resulting in a relatively fast reduction of genome size. For instance, coffee and tomato belong to the class Asteridae. Yet, since their divergence, the tomato lineage underwent a whole genome triplication[22]. Despite this, the tomato genome encodes only 36% more protein-coding genes than coffee, and has just one additional chromosome. Hence, chromosome number and gene content can serve to point to the existence of past polyploidization events, but are not precise indicators of the number or type of such events. Gene order (also known as synteny) is often used to assess past polyploidizations, generally by comparing the purported polyploid genome to a non-duplicated relative. However, this approach requires well-assembled genomes, and its power is limited for ancient events, as the signal is blurred by the accumulation of genome re- arrangements. Finally, phylogenomics provides an alternative approach to study past polyploidizations. In particular topological analysis of phylomes, which are complete collections of gene evolutionary histories, has served to uncover past polyploidization events[12,23–25]. Recently, phylome analysis was instrumental to distinguish between ancient auto- and allopolyploidization in yeast[26].

The olive tree (*Olea europaea* L.) is one of the most important fruit trees cultivated in the Mediterranean basin[27]. It belongs to the family Oleaceae (order Lamiales), which comprises other flowering plants such as the ash tree (*Fraxinus excelsior*) or jasmine (*Jasminum sambac*). The genome of *O. europaea* has a diploid size of 1.32 Gb distributed in 46 chromosomes (2n). Up to date, polyploids have been described within *O. europaea* as a recent polyploid series (2x, 4x, 6x) based on chromosome counting, flow cytometry and molecular markers[27]. However, little is known about ancient polyploidization in the olive tree and relatives. The complete genome sequence has recently been published[28], with analyses revealing an increased gene content as compared to other Lamiales. This highly suggested the existence of at least one past polyploidization event since the olive tree diverged from other sequenced Lamiales[28]. The recent sequencing of the genome of *F. excelsior* which also presents signs of a past WGD[29], further supports this hypothesis. Still,the exact number and nature of polyploidization events is yet to be resolved. To clarify this puzzle, we performed a phylogenomic analysis of the *O. europaea* genome.

## Results and discussion

### Gene order analysis indicates multiple polyploidizations in the Lamiales

A standard approach to confirm polyploidization relies on the finding of conserved syntenic paralogous blocks. Using COGE tools[30], we searched duplicated genomic regions in the olive genome. Our results revealed numerous such regions, which supports the existence of past polyploidization events (Additional file 1: Figure S1a). We then calculated the syntenic depth of the olive genome, which is a measure of the number of regions in the genome of interest that are syntenic to a given region in a reference non-duplicated genome (See Methods). As a reference we used *Coffea canephora*. This species belongs to the order Gentianales and, given the presence of duplications among sequenced Lamiales species, *C. canephora* is the closest non-duplicated reference genome[31]. As a control, we performed a similar analysis between *C. canephora* and *Sesamum indicum*, a Lamiales species known to have undergone a single WGD[32]. We also included *F. excelsior* (Oleaceae) in the comparison as the closest sequenced relative to the olive. Our analyses (Additional file 1: Figure S1b) revealed contrasting patterns between the three species. The *Sesamum-Coffea* comparsions showed a clear peak at depth 2, consistent with the reported WGD. In contrast, there was no such clear peak in the above mentioned *Olea-Coffea* or *Fraxinus-Coffea* comparisons, but rather a similarly high number of regions of depth 1 to 6, and 1 to 4, respectively. These results indicate the presence of multiple polyploidization events in the lineages leading to these species, and suggest that *O. europaea* may have undergone more such events than *F. excelsior*.

### The olive phylome

To elucidate the evolutionary history of *O. europaea* genes and compare it to that of related plants, we reconstructed the phylome[33] of this species and those of five other Lamiales (*F. excelsior, Mimulus guttatus, S. indicum, Utricularia gibba* and *Salvia miltiorrhiza*). These phylomes are available in PhylomeDB database[34] (see Additional file 2: Table S1 for details). We reconstructed the evolutionary relationships of the considered species using a concatenated approach with 215 widespread, single-copy orthologs (Fig. 1a), which yielded congruent results with previous analyses[35,36]. We scanned the trees to infer orthologs and paralogs, and date duplication events (see Methods). Using relative dating of gene duplications[37] we mapped them to the corresponding clades in the species tree. Functional analyses suggest that phosphatidylinositol activity, recognition of pollen, terpene activity, gibberellin metabolism and stress response are annotations enriched among genes duplicated in several of such periods (see Additional file 2: Table S2). We calculated the average duplication frequency for each marked node in Fig. 1b. Four internal branches showed increased duplication frequencies (nodes 2 to 5). In addition all terminal branches had high duplication frequencies and the two highest frequencies corresponded to the lineages of *U. gibba* (0.53 duplications/gene), for which two recent WGDs have been proposed[38], and to *O. europaea* (0.37). Altogether, these analyses indicate that the lineage leading to the olive tree shows three differentiated waves of massive gene duplications, one preceding the diversification of the sequenced Lamiales (node 4), another at the base of the family Oleaceae (node 5), and another one specific to the olive lineage.

**Fig. 1.**
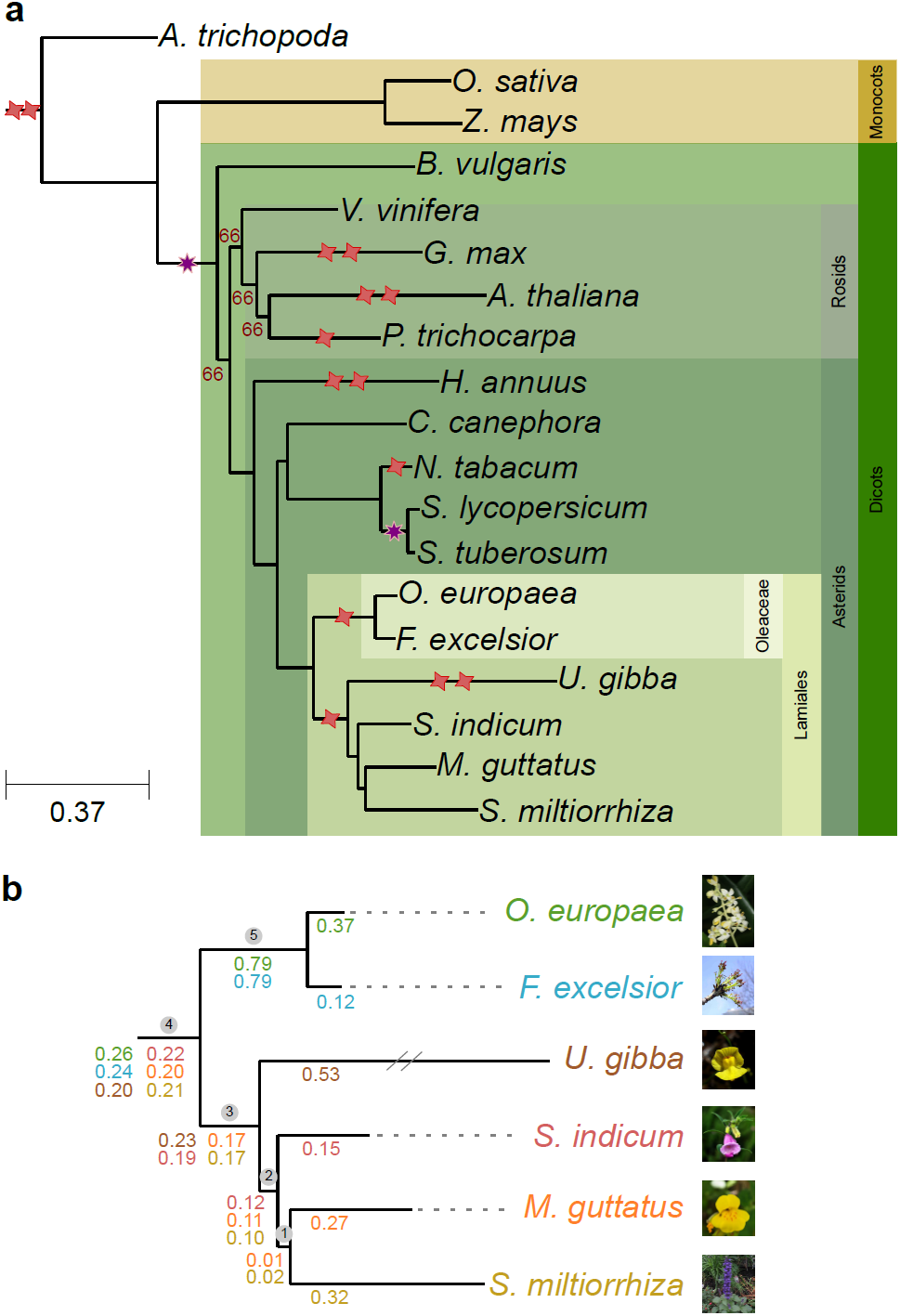
Species trees. (a) Evolutionary relationships between nineteen plants used in this study. All bootstrap values that are not shown in the graph, are maximal (100). Red stars represent WGD events, and purple stars represent whole genome triplication events, as described in the literature. **(b)** Zoom in to the Lamiales clade. Numbers in a circle on top of internal nodes represent the node names as referred to in the text, numbers below each branch are duplication frequencies calculated for each phylome. Each phylome and their corresponding duplication frequencies is colored differently: *O. europaea* - green, *F. excelsior* - light blue, *U. gibba* - brown, *S. indicum* - red, *M. guttatus* - orange, and *S. miltiorrhiza* - yellow.

### Phylogenetic analysis reveals an ancient allopolyploidization in Lamiales

We focused on the duplication peaks at the internal branches 2, 3 and 4 in Lamiales (Fig. 1b). A duplication event has been previously described within Lamiales[39], which could correspond to node 3 or node 4, depending on whether it is shared or not with Oleaceae. The peak at node 2, which has not previously been described, can be explained by the fact that the carnivorous plant *U. gibba*, despite the two recent WGDs, has a reduced genome resulting from massive gene loss[38]. Indeed for duplications that occurred at node 3, loss of all the duplicated paralogs in *U. gibba* would lead to mapping to node 2. Supporting such scenario is the finding that, when excluding orphan genes, only 51% of *S. indicum* genes have orthologs in *U. gibba* (see Additional file 3: Figure S2), as compared to 76% when comparing *S. indicum* to *M. guttatus* (see Additional file 3: Figure S2). To further test this scenario, we examined trees in the *S. indicum* phylome with node 2 duplications and counted how many of them included *U. gibba* homologs within the Lamiales clade. Only 20.7% of such trees fulfilled that pattern, further supporting that duplications mapped to node 2 mostly result from duplications occurred at node 3 followed by gene loss in *U. gibba*.

A similar scenario could explain duplications at node 3, if massive loss would have occurred in *O. europaea* and *F. excelsior*. Yet, these two species do not have reduced genomes (Additional file 3: Figure S2). In addition when scanning *S. indicum* phylome trees with either a duplication in node 2 or in node 3, homologs of *O. europaea* or *F. excelsior* could be found in 83.0% of them. Therefore, in this case, losses specific to Oleaceae cannot explain the duplication peak at node 3. This leads to the conclusion that at least two independent duplication events took place in the Lamiales: one corresponds to the previously described event[31,38] preceding the divergence of *M. guttatus* and *U. gibba* (node 3), and the other, congruent with a more ancestral event (node 4) preceding the divergence between Oleaceae and the other Lamiales species. To further confirm this newly discovered WGD (node 4), we performed a topological analysis on the 10,670 trees in the olive phylome presenting duplications at this node (see Methods), and assessed how many supported each of three possible topologies (see Fig. 2a): TA.- both paralogous lineages maintain gene copies in at least one species from both Oleaceae and the other Lamiales (non-Oleaceae) species; TB.- One of the paralogous lineages was lost in all non-Oleaceae Lamiales species; and TC.- One paralogous lineage was lost in all Oleaceae species. Surprisingly, many gene trees (77% in the *O. europaea* phylome) supported topology TB (see Fig. 2b). Equivalent analysis in the other Lamiales phylomes provided consistent results (Additional file 4: Figure S3).

**Fig. 2.**
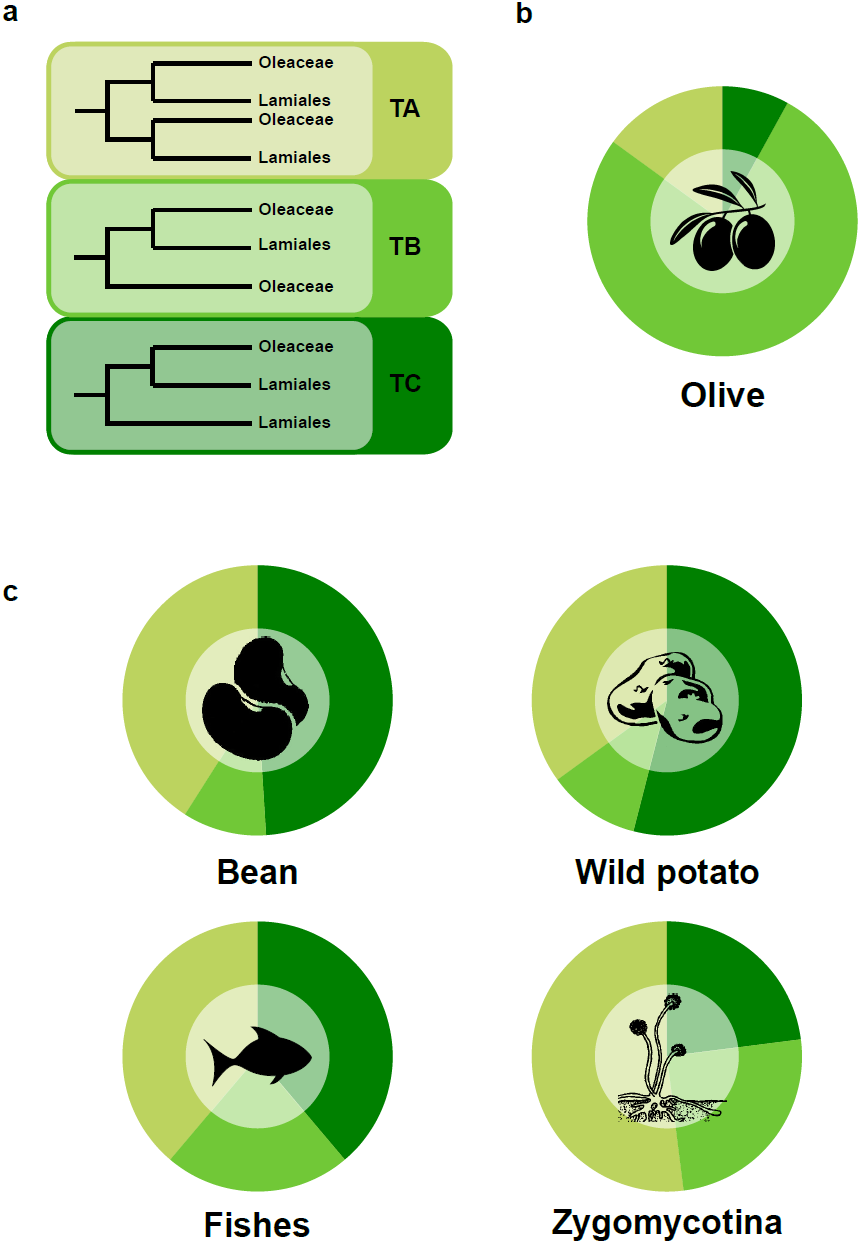
Topological analysis in olive and four other species. (a) Possible alternative topologies after the duplication concerning olive and the other Lamiales. **(b)** Percentage of trees that support each of the topologies shown in Fig. 2a in the olive phylome. **(c)** Percentage of trees that support each the different topologies for the phylomes of *Phaseolus vulgaris* (bean), *Solanum commersonii* (wild potato), *Scophthalmus maximus* (fishes), and *Rhizopus delemar* (Zygomycotina), taken from PhylomeDB. Like in Fig. 2a, TB indicates the loss of the paralogous side with the largest amount of species while TC indicates the loss of the paralogous side with the smallest amount of species.

We consider that a preponderance of topology TB is difficult to explain by a simple duplication and loss model. The imbalance in the number of species at the two sides of the node (two Oleaceae vs four non-Oleaceae Lamiales) means that, in scenarios involving gene losses, we expect a greater chance to observe topology TC than topology TB. This expected preponderance of TC was supported in analysis of other phylomes comprising WGD events at a node sub-tending imbalanced clades (see Fig. 2c). An alternative explanation for the preponderance of TB topology is the presence of an allopolyploidization at the base of Oleaceae. Indeed hybridization between an ancestor from a lineage that diverged before the Lamiales species included in our set and another species more closely related to the non- Oleaceae Lamiales would explain our observation[21,26] (see Additional file 5: Figure S4 for a detailed scenario).

### Increased phylogenetic resolution provided by transcriptomes uncovers allopolyploidization at the base of the tribe Oleeae

The ability to discern relative timing and type of past polyploidizations depends on the taxonomic sampling of the compared genomes. Unfortunately, at the time of starting this analysis the olive tree and *F. excelsior* were the only fully sequenced genomes from within the family Oleaceae. To increase the resolution of our analyses we included the transcriptomes of different Oleaceae species, whose genomes are not available: *Jasminum sambac*[40] and *Phillyrea angustifolia*[41]. The two species plus *F. excelsior* represent three important divergence points in the olive lineage. *P. angustifolia* belongs to the same subtribe (Oleinae), *F. excelsior* belongs to the same tribe (Oleeae) and *J. sambac* belongs to the same family (Oleaceae). In addition *J. sambac* has only 26 (2n) chromosomes, whereas the other three species have 46 chromosomes, which suggests that *J. sambac* likely experienced a lower number of polyploidizations. We thus expanded the olive phylome with these transcriptiomes (see Methods). We then selected two sets of trees: namely those including at least one sequence of each newly included species (set1: 20,705 trees) and those where a monophyletic clade contained the olive protein used as a seed in the phylogenetic reconstruction, and at least one sequence of each of the newly included species (set2: 11,352). Using the same approach described above we reconstructed the phylogeny of the expanded set of species (Fig. 3a), which was congruent with previous analyses based on plastid DNA[42]. Additionally we estimated their divergence times (see Methods and Additional file 6: Figure S5). The nodes (branches) in the new phylogeny were named from A to E (Fig. 3a), where E matched node 4 in the initial species tree (Fig. 1b). A new duplication profiling using set1 suggests three main duplication peaks in Oleaceae at nodes A, C, and D (see Additional file 7: Figure S6). The node at the base of the family Oleaceae (node D) is of similar density as the peak found at the base of the Lamiales (node E), which we already described as an allopolyploidization event that happened at the base of the Oleaceae family. Another peak at the base of the Oleeae tribe (node C) is higher than the previous two peaks, as could be expected of a more recent event. A third peak (node A) was still found specifically in *O. europaea*, indicating this duplication occurred after the divergence with *P. angustifolia*. Moreover, when duplication ratios are based on the more stringent set2 (see Additional file 7: Figure S6), ratios in nodes C and D are affected, while the rest remain with a similar density as in set1. The increased presence of proteins of *J. sambac* sister to the olive protein in gene trees in set2, can explain the increase in the ratio in node D, but not the decrease of the ratio in node C.

**Fig. 3.**
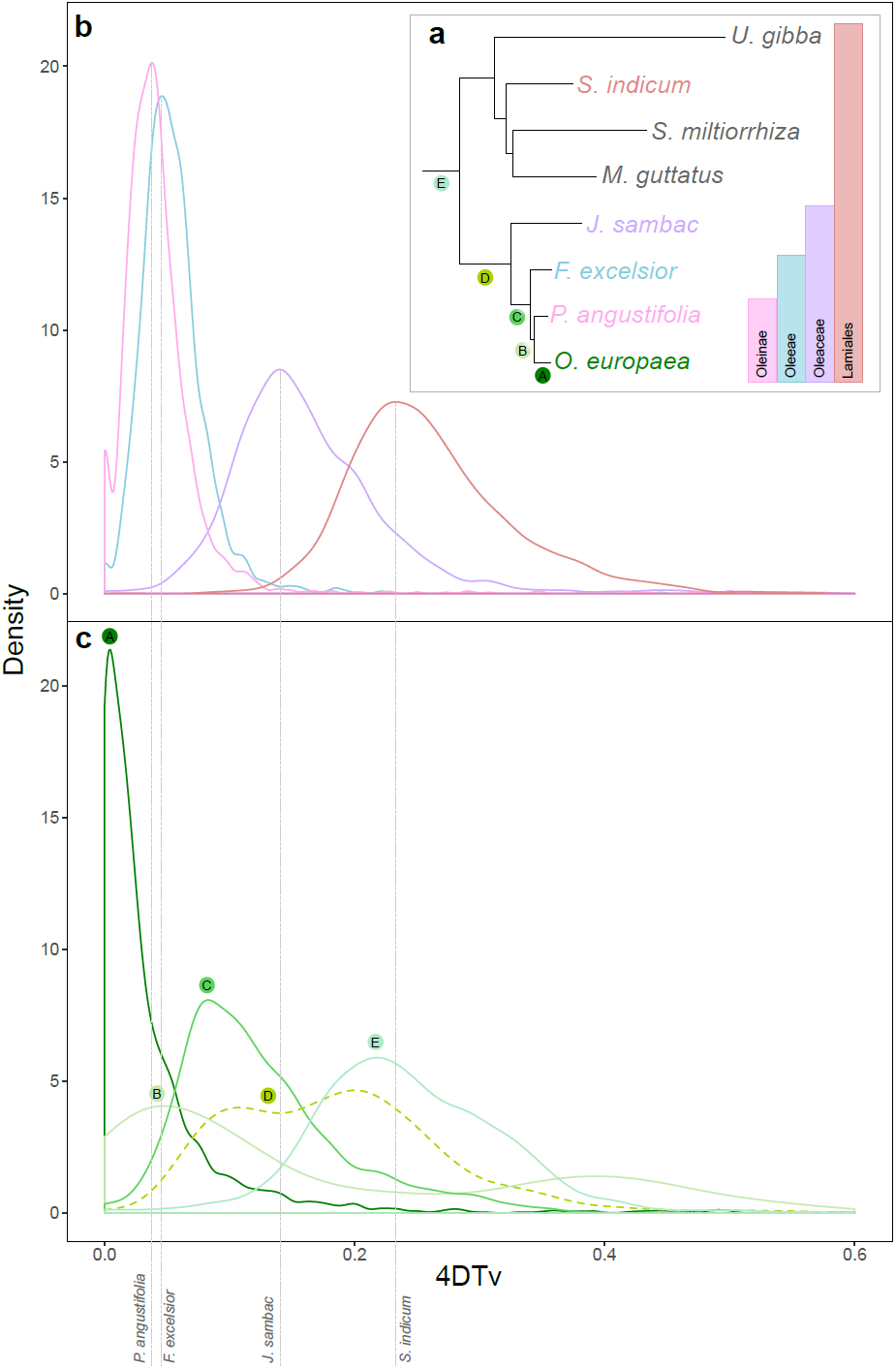
Species tree and 4DTv of the set1. (a) Species tree of the group of Lamiales including the additional two Oleaceae species, bars on the right show the taxonomic classification. Nodes where the 4DTv of the paralogous pairs were calculated are marked with letters (A to E) as referred to in the text and colored according each evolutionary age. The species used to calculate the 4DTv of orthologs pairs are shown in different colors **(b)** 4DTv of the orthologous pairs between *O. europaea* with *P. angustifolia, F. excelsior, J. sambac* and *S. indicum*. **(c)** 4DTv of the paralogous pairs of *O. europaea* at the marked nodes in the tree.

To obtain an independent assessment of the relative age of duplications, we plotted the ratio of transversions at fourfold degenerate sites (4DTv) for pairs of paralogs mapped at each of the branches in Fig. 3a, and compared these ratios with those of orthologous pairs found between *O. europaea* and the three other Oleaceae species plus *S. indicum* (see Fig. 3 and Additional file 8: Figure S7). The resulting patterns (Fig. 3) indicated overall congruence between topological dating and sequence divergence. The youngest peak comprised olive- specific duplications and followed the separation of olive and *P. angustifolia* of 10 MyA (see Additional file 6: Figure S5). The second wave of duplications appeared after the divergence of *J. sambac* and before the divergence of *F. excelsior*, at the base of the Oleeae tribe, which diverged between 14-20 MyA. Interestingly, duplications whose topology maps to two nodes appeared in this region of the 4DTv: those that map at node C after the divergence of *J. sambac* and a fraction of the duplications that happened before the divergence of *J. sambac* (node D). The most ancient duplication wave corresponds to the allopolyploidization event that we have previously described occurred between 33-72 MyA at the base of the Oleaceae family (node D). Of note this time frame includes the Cretaceous-Tertiary (KT) mass extinction event, around which many other plant polyploidization events have been predicted [11]. The fact that duplicatons whose topology map at node E are found in this region of the 4DTv, placed after the divergence of *S. indicum*, further supports the hybridization claim we propose. We also note that part of the duplications mapping at node D are found in this region. Altogether, these results confirm the presence of duplication three waves of duplications but also show that the duplications mapping at node D are divided in two peaks of sequence divergence as indicated by 4DTv plots. Node D duplications with 4DTv values found between the divergence of *S. indicum* and *J. sambac* can be explained as a result of the proposed allopolyploidization at the base of Oleaceae, either by the loss of non-Oleacae Lamiales species or by recombination where the non-Oleaceae Lamiales copy was over-written (Additional file 9: Figure S8). The other fraction of node D duplications with 4DTvs that map after the speciation of *J. sambac* are more difficult to explain, as in the trees they predate *J. sambac* divergence. This scenario is similar to the one we observe at the base of Oleaceae where topologically duplications are mapped at a different node than their age indicates. Therefore we propose that the Oleeae tribe was the result of a hybridization event with an ancestor in the lineage of *J. sambac* as one of the parents (Additional file 9: Figure S8). In 1945 Taylor proposed that the Oleaceae group with 23 chromosomes (Oleoideae) had an allopolyploid origin whose ancestors were two probably extinct lineages from a group related to *Jasminum* with chromosome numbers of 11 and 12 [43]. This scenario is further supported by the use of a more stringent filtering of the trees (set2). When at least one sequence of *J. sambac* is in the clade, then the duplication density at node D increases from 0.37 to 0.63 (Additional file 7: Figure S6). The use of a complete genome of *J. sambac* could confirm the allopolyploidization hypothesis at this point.

In order to confirm the two newly discovered allopolyploidization events with an alternative approach, we used GRAMPA [44], which relies on gene-tree species-tree reconciliation to discern between allo- or auto-polyploidization. We performed two different analyses. In the first we compared the allopolyploidization model versus the autopolyploidization model at the base of Lamiales (node E) (see Additional file 10: Figure S9a). We obtained lower parsimony scores for the allopolyploidization hypothesis (Additional file 2: Table S3), indicating a better match with the gene trees as compared to an autopolyploidization scenario. We performed the same analysis comparing the proposed allopolyploidization at the base of the Oleeae lineage (node C) with two different hypotheses that place an autopolyplodization at the base of the family Oleaceae and at the base of the tribe Oleeae, respectively (see Additional file 10: Figure S9b). The results once again supported allopolyploidization over each of the two autopolyplodization hypotheses. Finally, inspection of the phylome identified examples of gene trees that retained the duplications of the three polyploidization events, and whose topology is congruent with the proposed scenario (see Additional file 11: Figure S10 as an example). Re-analysis of the syntenic depth results uncovered over 800 homologous syntenic regions with a depth of 8 between coffee and olive (see Fig. 4 and Additional file 2: Table S4).

**Fig. 4.**
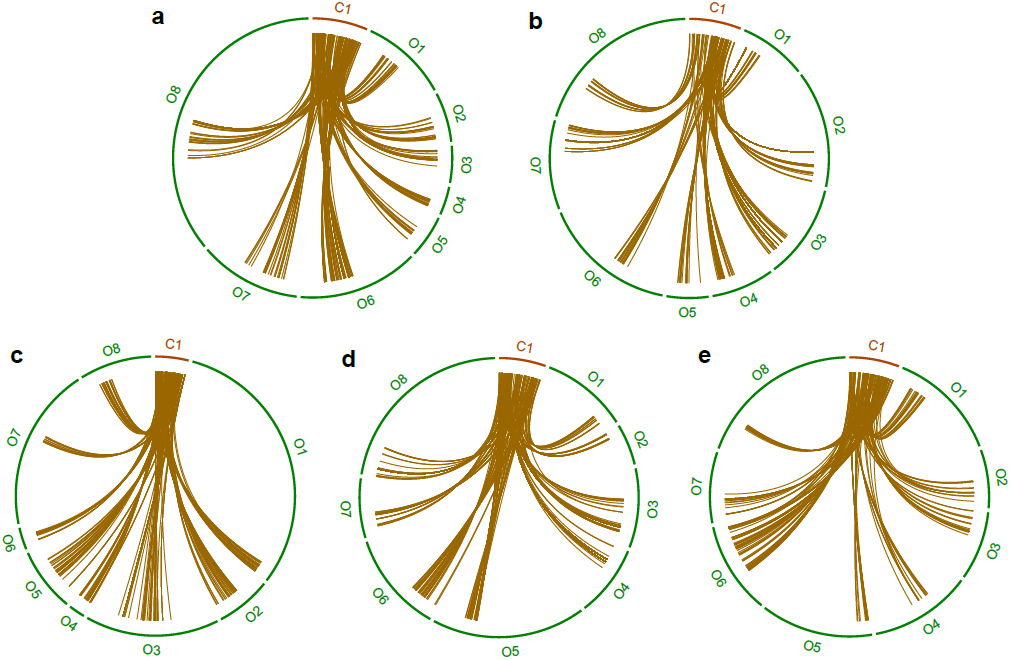
Example of five syntenic regions with a 1:8 relation between coffee and olive, as detected by GEvo. Exact regions can be found in Additional file 2: Table S4.

## Conclusions

Altogether our results underscore the power of phylogenomics to distinguish between allo- and auto-polyploidization. All our results indicate that the evolutionary history of olive comprises not only a species specific WGD, but also two older allopolyploidization events (Fig. 5). The most ancestral event occurred at the base of the family Oleaceae, where a non- Oleaceae Lamiales species could be involved as one of the parental species. Also this event is independent of the one described before for the group of non-Oleaceae Lamiales species. The second one at the base of the Oleeae tribe that seems to involve a species related to *Jasminum* as one of the partners. The third event is specific to *O. europaea* and, with the current set of sequenced species, we do not find phylogenetic support for an allopolyploidization scenario. However, increased taxonomic sampling may change this.

**Fig. 5.**
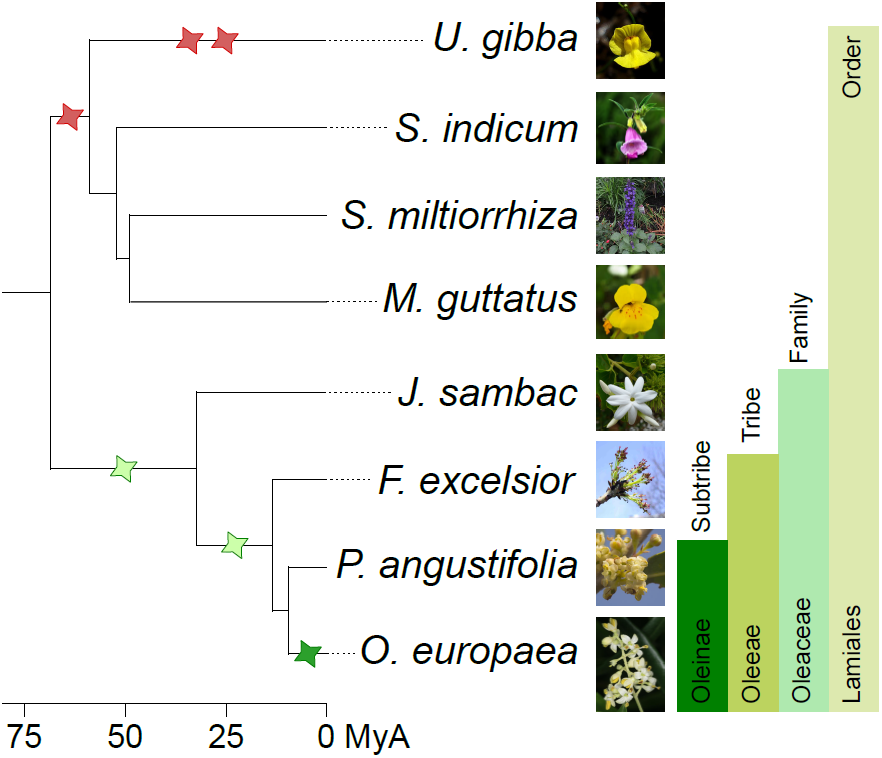
Species tree of the Lamiales clade showing the polyploidization events described in the literature (red stars) and in this analysis (green stars). The light green stars mark allopolyploidization events. Bars on the right show the taxonomic classification and the line in the bottom shows the divergence time in MyA.

## Methods

### Gene order analysis

The comparative genomic tools in the CoGe software package[30] (<u>https://genomevolution.org/coge/</u>) were used to analyse gene order in the genomes of olive and its relatives. First, synmap was used to compare the olive genome against itself using the Syntenic Path Assembly option[45] and removing scaffolds without conserved synteny (see Additional file 1: Figure S1). Then, we used SynFind to obtain the syntenic depth, the number of conserved syntenic regions between one query genome and a reference. We obtained this value for comparisons of the olive, *Fraxinus excelsior* and *Sesamum indicum* using *Coffea canephora* as reference (see Additional file 1: Figure S1). SynFind was also used to find regions with a 1:8 relationship between coffee and olive (see Fig. 4 and Additional file 2: Table S4).

### Phylome reconstruction

Six phylomes were reconstructed. In all cases an appropriate set of species was selected (see Additional file 2: Table S1) and the PhylomeDB automated pipeline was used to reconstruct a tree starting from each gene encoded in each one of the seed genomes[33]. This pipeline proceeds as follows: First a smith-waterman search is performed[46] and the resulting hits are filtered based on the e-value and the overlap between query and hit sequences (e-value threshold < 1e-05 and overlap > 0.5). The filtered results are then aligned using three different methods (MUSCLE v3.8, MAFFT v6.814b and KALIGN 2.04) used in forward and reverse orientation[47–50]. A consensus alignment is reconstructed from these alignments using M-coffee[51]. This consensus alignment is then trimmed twice, first using a consistency score (0.1667) and then using a gap threshold (0.1) as implemented in trimAl v1.4[52]. The resulting filtered alignment is subsequently used to reconstruct phylogenetic trees. In order to choose the best evolutionary model fitting each protein family, neighbor joining trees are reconstructed using BIONJ and their likelihoods are calculated using seven evolutionary models (JTT, WAG, MtREV, VT, LG, Blosum62, Dayhoff). The model best fitting the data according to the AIC criterion is then used to reconstruct a maximum likelihood tree with PhyML v3.1[53]. All trees and alignments are stored and can be downloaded or browsed in phylomeDB [54] (<u>http://phylomedb.org</u>) with the Phylome IDs 215, 216, 217, 218, 219, and 220.

### Incorporation of transcriptomic data in the olive phylome

Transcriptome data was downloaded from the ources indicated in their respective publications *Jasminum sambac*[40], and *Phillyrea angustifolia*[41]. In the case of *J. sambac*, where no protein prediction derived from the transcriptome was available, we obtained the longest ORF for each transcript. Only ORFs with a length of 100 aa or longer were kept, resulting in 20,952 ORFs in *J. sambac*. Transcriptomic data was introduced into each tree of the olive phylome using the following pipeline. First a similarity search using blastP was performed from the seed protein against a database that contained the two transcriptomes. Results were then filtered based on three thresholds: e-value < 1e-05, overlap between query and hit had to be at least of 0.3, and a sequence identity threshold > 40.0%. Hits that passed these filters were incorporated into the raw alignment of the phylome using MAFFT (v 7.222) (-add and -reorder options)[55]. Then trees were reconstructed using the resulting alignment and following the same procedure as described above. Once all trees were reconstructed, they were filtered to remove unreliably placed transcriptome sequences. Phylomes tend to be highly redundant, specially when the seed genome contains many duplications, as is the case for the olive genome. Therefore, the same transcriptomic sequence is likely inserted in many trees. For each inserted transcript, we checked whether the sister sequences of each inserted transcript overlapped. If such overlap did not exist the transcript was deemed unreliable and removed from the tree. This filtered set was then filtered once more to select trees that contained at least one transcript for each of the two new species (set1). Finally set1 was filtered again to keep only trees that contained a monophyletic clade including the four *Oleaceae* species (set2).

### Species tree reconstruction

A species tree was reconstructed using data from the olive phylome. Each tree reconstructed for this phylome was first pruned so that species specific duplications were deleted from the tree, keeping only one sequence as representative of the duplicated group. Once trees were pruned, only those trees that contained one sequence for each of the 19 species included in the phylome were selected. 215 such trees were found. The clean alignments used to reconstruct these trees were concatenated and a species tree was reconstructed using the model of amino acids substitution LG implemented in PhyML v3.1[53] with 100 bootstrap replicates. In addition, a second species tree was reconstructed using a super-tree approach with the tool duptree[56]. In this case all trees in the olive phylome were used for the tree reconstruction. A third species tree was reconstructed after the inclusion of the transcriptomic data into the olive phylome. From the initial set of genes chosen to reconstruct the first species tree, a subset was chosen to reconstruct the extended species tree. This subset included only genes that incorporated at least one of the three species with a transcriptome. This final tree was reconstructed using 112 gene alignments using the same methodology as described above.

### Detection and mapping of orthologs and paralogs

Orthologs and paralogs were detected using the species overlap method[25] as implemented in ETE v3.0[57]. Species specific duplications (expansions) are computed as duplications that map only to one species, in our case always the species from which the phylome was started. In order to reduce the redundancy in the prediction of species specific expansions a clustering is performed in which expansions that overlap in more than 50% of their sequences are fused together.

Predicted duplication nodes are then mapped to the species tree under the assumption that the duplication happened at the common ancestor of all the species included in the node, as described by Huerta-Cepas and Gabaldón[37]. Duplication frequencies at each node in the species tree are calculated by dividing the number of duplications mapped to a given node in the species tree by all the trees that contain that node. In all cases duplication frequencies are calculated excluding trees that contained large species specific expansions (expansions that contained more than 5 members).

### GO term enrichment

GO terms were assigned to the olive proteome using interproscan[58] and the annotation of orthologs from the phylomeDB database[54]. Phylome annotations were transferred to the olive proteome using one-to-one and one-to-many orthologs. GO term enrichment of proteins duplicated at the different species-specific expansions and duplication peaks was calculated using FatiGO[59].

### Topological analysis

A topological analysis was performed using ETE v3.0[57] to test whether a duplication event happened at the base of *Lamiales* and determine which species were involved. We searched how many trees supported each of the following topologies: the complete topology where at least one *Oleaceae* and at least one other non-*Oleaceae Lamiales* are found at both sides of the duplication (topology TA), a partial topology where all non-*Oleaceae Lamiales* species have been lost in one side of the duplication (topology TB), or another partial topology where the *Oleaceae* sequences have been lost at one side of the duplication (topology TC) (see Fig. 2a). The analysis was then repeated in different previously reconstructed phylomes that contained ancient whole genome duplications where there was an imbalance of species at either side of the duplication. The phylomes selected were those of the plants *Phaseolus vulgaris*[60] (Phylome ID 8) and *Solanum commersonii*[61] (Phylome ID 147), the fish *Scophthalmus maximus* [62](Phylome ID 18), and the fungi *Rhizopus delemar*[23] (Phylome ID 252). Each of those phylomes contains an old WGD where at one side of the duplication there are less species than at the other one. We checked the proportion of trees that supported each topology. Like with the *Oleaceae* example, topology TA’ conserves at least one member of each group, topology TB’ has lost all the species of the large group at one side of the duplication while TC’ has lost all the species of the small group at one side of the duplication (see Fig. 2d).

We used GRAMPA[44] (Spring 2016 version) to assess five different hypothesis (see Additional file 11: Figure S10) using the two sets of trees that contained transcriptomic data. This tool uses reconciliation in order to compute the support between a set of trees and a proposed allopolyploidization or autopolyploidization event. Though it is limited to detecting one single event at a time. During its calculation, GRAMPA discards single gene trees that have too many possibilities when reconciling them to the species tree. The trees discarded can vary depending on the species tree hypothesis. Therefore, in order to fairly compare the parsimony scores obtained, we recalculated them based on the trees used in all the hypotheses. We performed two different analyses. In the first we compared the allopolyploidization model versus the autopolyploidization at the base of *Lamiales* (see Additional file 11: Figure S10a). In the second we compared the allopolyploidization that led to the *Oleeae* lineage with two different hypotheses that place an autopolyploidization at the base of *Oleaceae* family and at the base of *Oleeae* tribe respectively (see Additional file 11: Figure S10b). Results can be found in Additional file 2: Table S3.

### Transversion rate at fourfold degenerate sites (4DTv)

The 4DTv distribution was used to estimate speciation and polyploidization events. In order to obtain the gene pairs we used the species tree that included the transcriptomic data. We calculated the 4DTv values for the orthologous gene pairs between *O. europaea* with *J. sambac, F. excelsior, P. angustifolia*, and *S. indicum*. We also calculated the 4DTv for each paralogous gene pair of olive that maps at each evolutionary age.

### Divergence times

Divergence times were calculated using r8s-PL 1.81 [63]. Four nodes were taken as calibration points. The divergence time of these nodes were obtained from the TimeTree database [64]: *Mimulus guttatus* - *Arabidopsis thaliana* (117 MYA), *Sesamum indicum* - *Solanum lycopersicum* (84 MYA), *Glycine max* - *Arabidopsis thaliana* (106 MYA), *Zea mays*- *Solanum lycopersicum* (160 MYA). Cross-validation was performed to choose the smoothing parameter.

## Declarations

### Author’s contributions

IJ and MMH performed bioinformatics analysis. IJ, MMH, and TG analyzed the results. TG and PV supervised the study. All authors wrote, read and approved the manuscript.

## Acknowledgments

TG group acknowledges support from the Spanish Ministry of Economy and Competitiveness grants, ‘Centro de Excelencia Severo Ochoa 2013-2017’ SEV-2012-0208, and BFU2015-67107 cofounded by European Regional Development Fund (ERDF); from the European Union and ERC Seventh Framework Programme (FP7/2007-2013) under grant agreements FP7-PEOPLE-2013-ITN-606786 “ImResFun” and ERC-2012-StG-310325; from the Catalan Research Agency (AGAUR) SGR857, and grant from the European Union’s Horizon 2020 research and innovation programme under the Marie Sklodowska-Curie grant agreement No H2020-MSCA-ITN-2014-642095; TG and PV acknowledge support from Banco Santander for the olive genome sequencing project; IJ was supported in part by a grant from the Peruvian Ministry of Education: ‘Beca Presidente de la República’ (2013-III).

## Availability of data and materials

All data generated or analyzed during this study are included in this published article and its supplementary information files, or is available upon request.

## Additional files

**Additional file 1: Figure S1. Results obtained with the GOGE package. (a)** Image of a mapping of *O. europaea* against itself as shown by Synmap. **(b)** Syntenic depth as calculated by SynFind.

**Additional file 2: Tables:**

**Table S1. List of species included in the reconstruction of the six phylomes used in this study.** Columns indicate, in this order, the species code for each species, the species name, the source for the protein and the coding DNA sequences, and the phylome in which the species was used (*O. europaea*-215, *F. excelsior*-216, *M. guttatus*-217, *S. indicum*-218, *U. gibba*-219, *S. miltiorrhiza*-220).

**Table S2. List of the GO terms enriched in the expanded protein families and at each evolutionary period as described in Fig. 1b.** First column shows the GO term, the second the term level, the third, the p-value, and the fourth, the term name.

**Table S3. List of parsimony scores for each of the different hypothesis and considering the two sets of trees with EST data.** Nodes are named as shown in Fig. 3.

**Table S4. Syntenic regions between coffee and olive used in the Fig. 4.** In the first column we can see the letter of the graph. The second and the sixth columns show the scaffold names used in the graph (names starting with “C” are for coffee and “O” are for olive). The third and the seventh columns show the scaffold names of the genome in coffee and olive, respectively. The fourth and the fifth columns show the start and the end of the region in coffee. The eighth and the ninth columns show the start and the end of the syntenic region in olive.

**Additional file 3: Figure S2. Heatmap showing the percentage of orthologous proteins in comparison to each Lamiales species included in this analysis.**

**Additional file 4: Figure S3. Pie-charts representing the distribution of trees supporting each of the topologies as shown in Fig. 2.**

**Additional file 5: Figure S4: Exact topologies expected to find in a scenario of autopolyploidization and one of allopolyploidization.**

**Additional file 6: Figure S5. Chronogram depicting the evolution of the plants included in the phylome. Green dots represent selected calibration points in MyA.**

**Additional file 7: Figure S6. Species tree of the Lamiales order, including *P. angustifolia, F. excelsior* and *J. sambac***. The duplication rates are shown in red for set1 and in blue for set2. The grey circles show the node name and the bars on the right, the taxonomic classification.

**Additional file 8: Figure S7. Species tree and 4DTv of the set2. (a)** species tree of the group of Lamiales including the three Oleaceae species. Nodes where the 4DTv of the paralogous pairs were calculated are marked with letters (A to E) as referred to in the text and coloured according each evolutionary age. The species used to calculate the 4DTv of orthologous pairs are shown in different colours. The bars on the right show the taxonomic classification. **(b)** 4DTv of the orthologous pairs between *O. europaea* with *P. angustifolia, F.excelsior, J. sambac* and *S. indicum*. **(c)** 4DTv of the paralogous pairs of *O. europaea* at the marked nodes in the tree.

**Additional file 9: Figure S8. Schematic explanation of the 4DTv density at node D in Fig. 3c. (a)** representation of the two allopolyploidization events and the potential parentals.**(b)** scheme of a gene tree where the protein of *J. sambac* map after the divergence of this species. **(c)** scheme of a gene tree where the non-Oleaceae Lamiales proteins are lost. **(c)** 4DTv of the paralogs at nodes C, D, and E. The dotted lines mark the divergence time between olive - *J. Sambac* and olive - *S. indicum*.

**Additional file 10:** Figure S9. Phylogenetic trees representing the comparisons done for GRAMPA. In all cases branches painted in green and orange represent the species that the polyploidy has affected. **(a)** The trees represent the hypothesis of an allopolyploidization versus an autopolyploidization at the base of Lamiales. **(b)** These trees represent the hypothesis of an allopolyploidization versus a two models of autopolyploidization.

**Additional file 11: Figure S10. Example gene tree that shows the three events we have described in olive:** the species specific duplication and the two allopolyploidizations. The whole genome duplication previously described in non-Oleaceae Lamiales and the species specific duplications in *U. gibba* can also be seen.

